# Divergent spatiotemporal integration of whole-field visual motion in medaka and zebrafish larvae

**DOI:** 10.1101/2025.08.22.671687

**Authors:** Yasuko Isoe, Yasmine Fatima Mabene, Marie-Abèle Bind, Florian Engert

## Abstract

Cross-species comparisons offer leverage for identifying conserved and divergent neural computations underlying innate behavior. Visual motion integration is a fundamental operation that stabilizes position relative to the moving environment and supports object tracking, yet how its underlying algorithms vary across closely related vertebrate brains remains poorly understood. We investigated how zebrafish (*Danio rerio*) and medaka (*Oryzias latipes*) larvae implement visual motion integration using distinct spatiotemporal filters that trade speed for persistence through separable control modules. Using controlled whole-field motion stimuli, we found that medaka pool motion signals over visual fields nearly twice as large as those of zebrafish and exhibit enhanced weighting of peripheral inputs, whereas zebrafish rely more strongly on motion signals directly beneath the body. Temporally, zebrafish respond robustly to motion signals with lifetimes as short as 100 ms, whereas medaka require stimulus durations exceeding one second and maintain motion-driven activity for several seconds after stimulus offset. Decomposition of turning behavior revealed separable control modules for large and small corrective maneuvers, with species differences arising primarily from prolonged temporal integration in medaka small-turn control. Together, these differences reveal species-specific tuning of spatial kernels and temporal filters underlying visuomotor control. Our results demonstrate how alterations in basic computational motifs, spatiotemporal pooling, gain, and persistence, can generate divergent visuomotor strategies across closely related vertebrate brains.

**Significance Statement:** Animals rely on visual motion to stabilize positions and interact with dynamic environments, yet how these computations vary across related species remains unclear. By comparing larval zebrafish and medaka, we show that visually similar vertebrates implement motion integration using distinct spatiotemporal strategies. Medaka integrate motion over larger visual fields and retain motion signals for seconds, whereas zebrafish favor rapid, spatially restricted integration. These differences arise from separable control modules governing fine and large motor adjustments. Our results reveal how small changes in core computational motifs—pooling, gain, and persistence—can generate divergent sensorimotor strategies across evolution.

## Introduction

Cross-species comparisons provide a powerful framework for identifying both conserved and divergent neural computations underlying behavior [1–3]. Even when animal species display similar behavioral repertoires, the underlying sensory processing and sensorimotor transformations can differ substantially, reflecting adaptation to distinct ecological and environmental demands. Understanding how such computational strategies vary across species offers insight into the flexibility and evolutionary tuning of neural circuits.

Visual motion integration is a fundamental computation that enables animals to stabilize their position relative to the environment, extract meaningful cues from dynamic visual scenes, and guide locomotor decisions [4]. One of the most prominent behaviors relying on visual motion integration is the optomotor response (OMR), a reflexive orienting behavior elicited by whole-field visual motion [5–7]. During OMR, animals adjust their orientation and movement to counteract perceived drift of the visual scene, thereby maintaining positional stability in moving environments such as flowing water. Although OMR is often described as a simple reflex, it relies on sophisticated processing of motion signals distributed across space and time [8].

Motion integration involves at least two core computational dimensions. Spatial integration pools motion signals across the visual field, combining local motion estimates while resolving conflicts between competing motion directions. Many species exhibit region-specific sensitivities to whole-field motion, reflecting ecological constraints such as habitat structure or typical patterns of self-motion. Temporal integration, in turn, governs how motion information is accumulated and retained over time. Short temporal filters allow rapid responses to transient motion cues, whereas longer integration times promote stability by suppressing brief fluctuations and retaining motion information after stimulus offset. Together, spatial pooling and temporal filtering determine how motion signals are converted into appropriate motor commands.

Motion detection engages computations across two core dimensions: space and time. Spatial integration pools motion information across the visual field, by summation over coherent motion and by performing antagonistic computations for opposing patterns [9, 10]. Many species also show region-specific whole field motion sensitivities that reflect specialized ecological demands [11–13]. Temporal integration enables the accumulation and retention of motion evidence, supported by neural mechanisms capable of sustaining activity across hundreds of milliseconds or longer [14, 15]. In natural settings, temporal statistics of motion help distinguish noise from meaningful cues: brief, stochastic fluctuations likely represent environmental variability, whereas long-lifetime motion typically signals objects or self-motion [16]. Thus, species evolving under different ecological demands are expected to diverge in the structure of their spatial kernels and temporal filters.

Despise extensive characterization of motion processing within individual speces, comparatively little is known about how these spatiotemporal integration strategies differ across closely related vertebrates. Most studies examine motion integration in isolation, focusing on a single model organism and treating temporal persistence or spatial extent as fixed properties rather than adjustable computational variables. As a result, it remains unclear how evolution tunes the balance between responsiveness and stability, or how small changes in core filtering principles can give rise to distinct sensorimotor strategies.

Larval teleost fish larvae offer an ideal system for addressing these questions. Zebrafish (*Danio rerio*) and medaka (*Oryzias latipes*) diverged ∼200 million years ago, yet exhibit broadly similar morphologies during early development. In zebrafish larvae, moving whole field visual stimuli elicit robust OMR [17, 18] and the underlying circuitry—from retinal processing to hindbrain sensorimotor pathways—has been extensively characterized [9, 14]. In contrast, although medaka OMR has been documented in juveniles and adults and used to probe visual feature sensitivity [19, 20, 21], the computational rules governing motion integration in medaka larvae remain largely unexplored. This combination of behavioral similarity and phylogenetic divergence makes zebrafish and medaka a powerful comparative pair for probing the evolution of motion-processing algorithms.

Here, we quantitatively compare optomotor behaviors in zebrafish and medaka larvae using controlled whole-field motion stimuli that systematically vary spatial extent and temporal structure. By analyzing turning behavior across multiple stimulus paradigms, we infer species-specific spatial kernels and temporal filters underlying motion integration. Decomposition of turning responses reveals separable control modules for large and small corrective maneuvers, allowing us to isolate the computational components that differ between species. Finally, we introduce a mechanistic but phenomenological model that captures key differences in gain and temporal persistence and reproduces species-specific response dynamics. Together, our results demonstrate that closely related vertebrates implement visual motion integration using distinct spatiotemporal strategies, revealing how modest changes in core computational motifs can generate divergent visuomotor control strategies.

## Results

### Cross-species comparison in optomotor response (OMR) in teleost larvae

We leverage cross-species comparison as a natural perturbation to the sensorimotor system, and here we compared larvae of two teleost species, zebrafish and medaka (**Fig 1 a, b**). While both species can be found in rice field environments, they share the same behavioral repertoire, however they diverged ∼200 million years ago [22]. Given that adapting to environmental visual cues is crucial for larval survival, this study aimed to determine whether these species have evolved different behavioral strategies for responding to whole-field motion stimuli.

**Figure 1.**
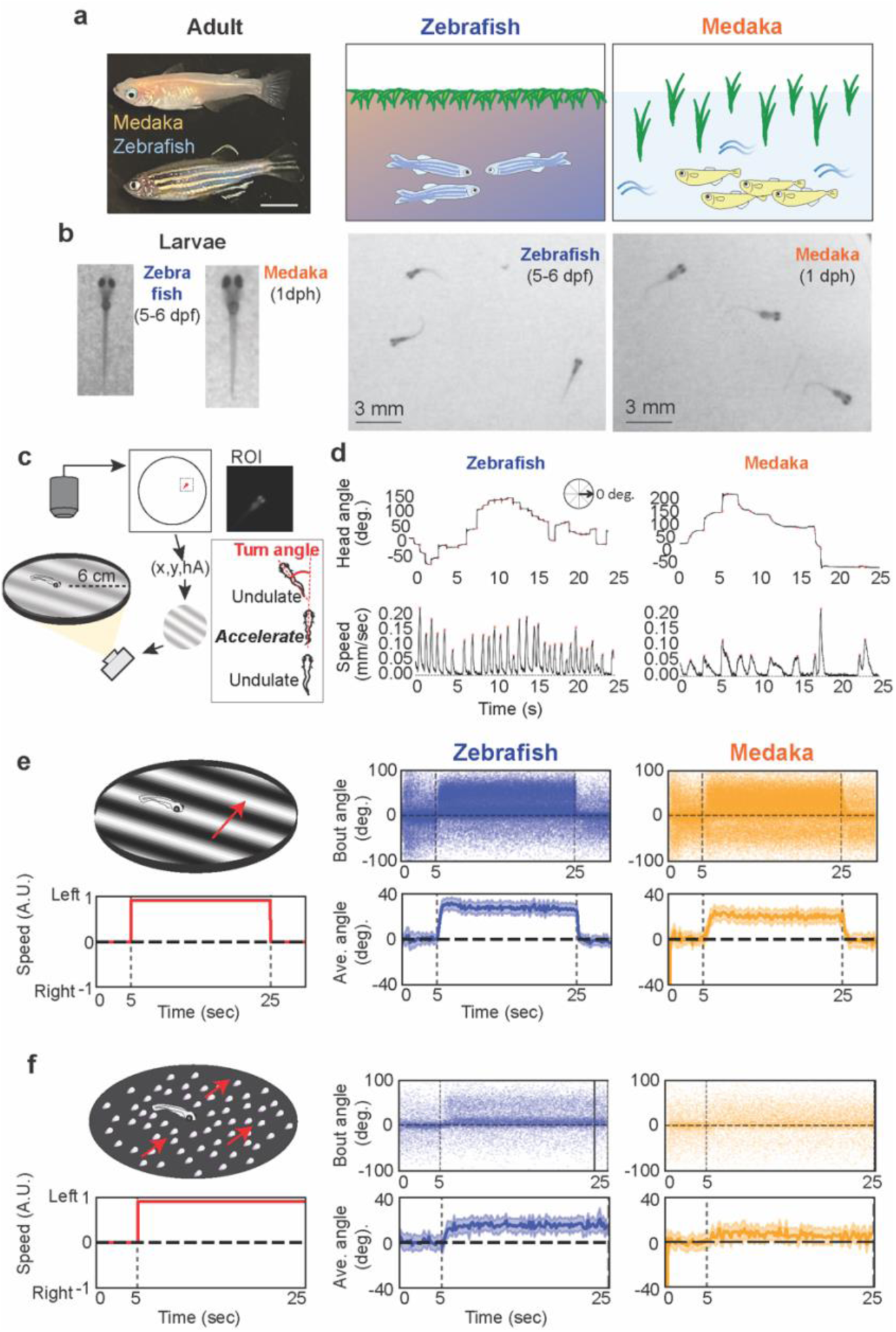
Comparative study of optomotor response (OMR) in zebrafish and medaka larvae. (a) Zebrafish and medaka species comparison. Adult fish (left; top: medaka, bottom: zebrafish) and schematic drawings are shown. Scale bar: 1 cm. (b) Larval specimens used in this study. Larvae at 5-6 days post-fertilization (dpf) for zebrafish and 1 day post-hatch (dph) for medaka were examined. Scale bar: 3 mm. (c) Free-swimming behavioral experiment rig. Individual larvae were placed in a 6 cm-radius circular arena. A high-speed camera above the dish took the whole image of the behavioral rig at 90 fps. The center of mass position (x, y) and head angle (hA) of the focal larvae are recorded. Visual stimuli were generated based on these parameters and projected from beneath the arena. (d) Swimming dynamics of zebrafish (left) and medaka (right) larvae. Zebrafish exhibit discrete swimming bouts alternating with interbout periods, whereas medaka demonstrate continuous undulatory swimming with rhythmic acceleration phases. Turn angles were quantified as the deviation in head angle before and after acceleration time points, as illustrated in the bottom right inset of (c). (e) Optomotor response to whole-field sine-grating stimuli (top left). The red trace represents the temporal stimulus pattern (bottom left), where positive values indicate leftward motion and negative values indicate rightward motion. Each trial consisted of 5 seconds open-loop converging sine-grating baseline, followed by 20 seconds closed-loop sine-grating stimuli, and 5 seconds static grating. Turn angles for all swimming bouts are plotted (right, top row; blue: zebrafish, orange: medaka), with averaged responses with shades of the standard errors of the means (SEMs) shown below (right, bottom row) (zebrafish: n=53, medaka: n=42). (f) Optomotor response to whole-field random dot motion stimuli (top left). Protocol identical to (e) 5 seconds open-loop converging sine-grating baseline, followed by 20 seconds closed-loop random dot motion stimuli (bottom left). Individual bout turn angles (top row) and averaged responses with shades of SEMs (bottom row) are shown (zebrafish: n=21, medaka: n=32).

To enable meaningful cross-species behavioral comparisons, we first identified comparable developmental stages. Based on literature describing organ development [22] and our observations of locomotor ontogeny (**S1 Fig a**), we determined that 5-6 days post-fertilization (dpf) zebrafish and 0-1 days post-hatch (dph) medaka exhibit similar swimming maturity. Specifically, 5–6 dpf zebrafish and 7–8 dpf (0–1 dph) medaka demonstrated comparable regular swimming patterns and were therefore used for all behavioral experiments. We utilized a behavioral assay in which visual stimuli were projected from below the experimental arena while fish swimming behavior was recorded from above (**Fig 1c**). To quantitatively assess optomotor responses, we measured turn angles as the deviation in head orientation before and after movement bouts (**Fig 1d**). While zebrafish swim in discrete bouts making turn angle calculation straightforward, medaka exhibit continuous tail undulation but also display discrete acceleration bouts during which head angle changes occur, enabling comparable measurements across species. As a complementary metric, we calculated the fraction of turns aligned with stimulus direction as the ‘correct response fraction’ [14, 23, 24] (**S1 Fig b**).

We characterized optomotor responses using two stimulus types: sine-wave gratings (**Fig 1e**, **S1 Fig. b, S1 Movie**) and flickering dot motion (**Fig 1f**, **S1 Fig. b**). Each trial began with 5 seconds of open-loop converging motion to center the fish, followed by closed-loop whole-field motion stimuli that were locked to fish position and head angle. Both species exhibited reliable and consistent directed turning toward the direction of moving whole-field stimuli, as evidenced by changes in average turning angular magnitudes (**Fig 1e, f**) and correct response fractions (**S1 Fig b**), but zebrafish generally exhibited more vigorous optomotor responses compared to medaka, as measured by both turn angle magnitude (**Fig 1e, f**) and response consistency (**S1 Fig b**). These initial observations suggested species-specific differences in visual-motor processing that warranted further investigation.

### Local and global spatial integration

From our initial observations, we noticed that larval turning behavior varied systematically with distance from the arena wall. Specifically, proximity to the wall reduced the ability to execute large-angle turns in the stimulus direction, suggesting that the wall acts as a visual ‘distractor’ (**Fig 2a, b** left**, c** top**, d** solid lines**, e** top). Average turn angles were significantly higher when larvae were positioned near the center of the arena in both zebrafish (blue) and medaka (orange), and decreased progressively as fish moved closer to the wall (**Fig 2 d, e**, solid lines). This pattern suggests that turning behavior is driven by the integrated motion energy within the fish’s visual field, with the black arena wall apparently inhibiting turning responses by occupying visual space with non-moving dark regions. Importantly, this wall-proximity effect was more pronounced in medaka than in zebrafish, indicating species differences in spatial integration.

**Figure 2.**
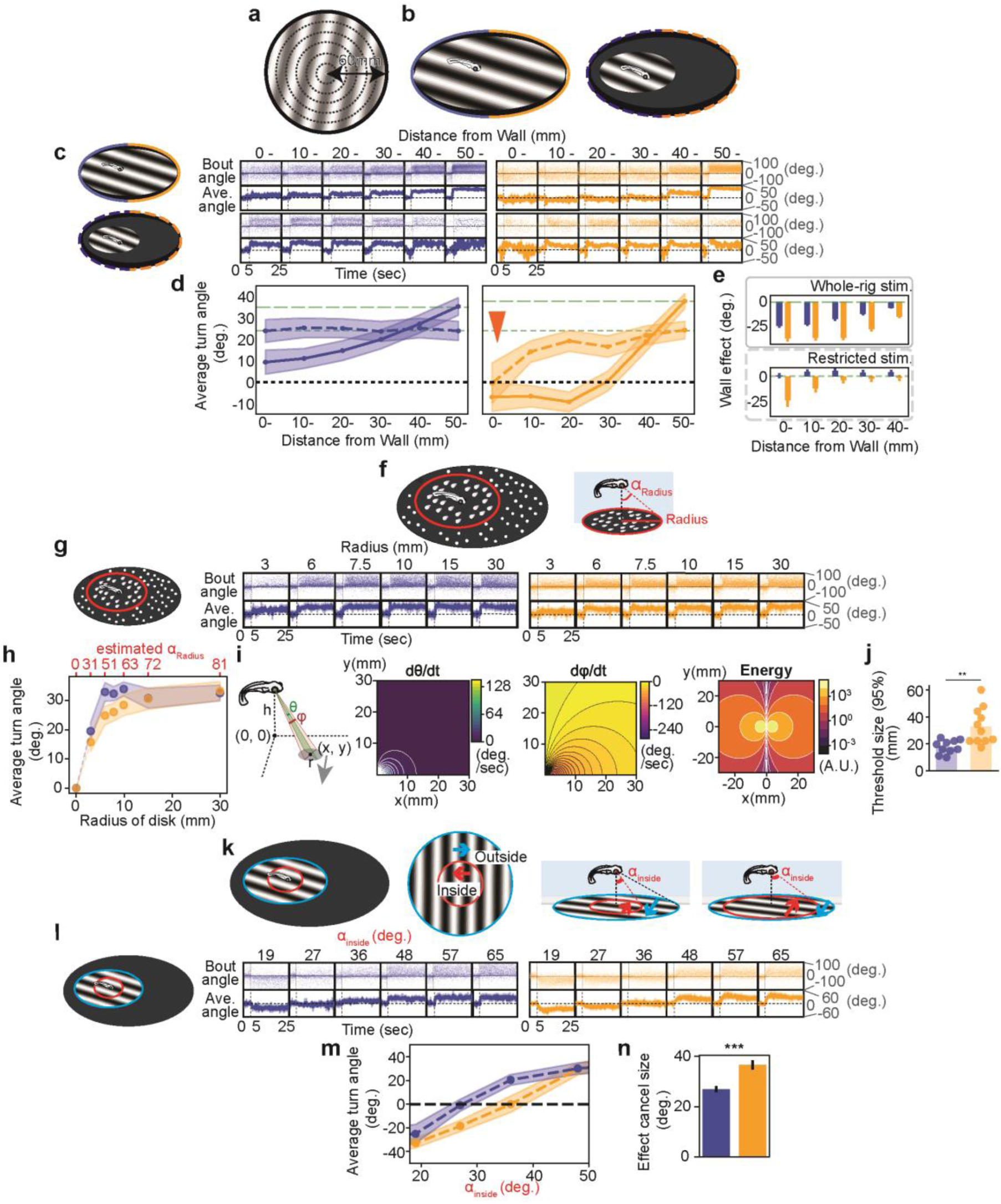
Spatial integration of the whole-field motion stimuli in teleost larvae. (a) Schematic of experimental arena. Turns were categorized based on distance from the wall where each turn occurred. (b) Sine-grating stimulus configurations: whole-field stimuli (left) or restricted area around the fish (right, 3 cm radius). (c) Turn responses at different distances from the wall. Raw angular data (top) and averaged turn angles across spatial bins (bottom) during trials with 5 s open-loop converging stimuli followed by 20 s closed-loop stimuli. Top rows: whole-field sine-grating stimuli (solid frames) (zebrafish n=42, medaka n=53). Bottom rows: restricted-area stimuli (dotted frames) (zebrafish n=33, medaka n=31). Blue: zebrafish, orange: medaka. (d) Averaged turn angles during closed-loop stimuli from (c). Solid lines: whole-field stimuli; dotted lines: restricted stimuli. Green dotted lines indicate the values of the responses at the wall-nearest area in the whole-rig stimuli (longer dot) and restricted stimuli (shorter dot) condition. Mean turning angles (± SEM) across six spatial bins from wall (bin 0) to center (bin 50). Statistical significance determined by Bonferroni-corrected t-tests following significant stimulus × spatial bin interaction in two-way mixed ANOVA (*p < 0.05, p < 0.01, p < 0.001). Orange arrowhead indicates persistent wall effect. (e) “Wall effects” calculated as deviation between each area and wall-nearest area. With whole-field stimuli, wall effects exist significantly in both species; with restricted stimuli, only medaka show significant wall effects. (f) Moving dots presented in disks of varying radii beneath fish, with stationary dots in surrounding areas. Visual occupying angle (α_disk_) estimated for fish swimming 5 mm above sandblasted dish bottom (See Supplementary figure 2d). (g) Turn responses to different motion disk sizes. Raw angular data (top) and averaged angles (bottom) (zebrafish n=10, medaka n=12). (h) Averaged turn angles during 5-25 s across motion disk sizes. Shaded areas: standard errors. Dotted lines: fitted curves based on motion energy calculations. (i) Motion energy calculation for larvae at height h. Each dot at position (x, y) occupies a visual field angle (θ, φ). For x-axis motion, angle changes depend on position (left: dθ/dt, middle: dφ/dt), and motion energy (dθ/dt × dφ/dt) varies by position (right). (j) Larger disks produce greater turning angles, but zebrafish reach 95% maximum response at smaller sizes than medaka (zebrafish: 17.8 ± 4.8 degrees, medaka: 32.8 ± 12.9 degrees; Welch’s t-test, p=0.0044). (k) Inconsistent motion stimuli with opposing directions in inner and outer regions. Response measured across different inner disk angular sizes. (l) Turn responses to varying inner disk sizes. Raw bout data (top) and averaged angles (bottom) (zebrafish n=6, medaka n=11). (m) Average turn responses versus inner disk size. Black dotted line indicates “effect cancel size” where inner and outer motions balance. (n) Effect cancellation size comparison. Medaka showed significantly larger cancellation angular size than zebrafish (zebrafish: 27.0 ± 1.2 degrees, medaka: 36.6 ± 2.0 degrees; Student’s t-test, p=0.00099).

Notably, medaka displayed a tendency to execute turns opposite to the stimulus direction when swimming close to the wall (<30 mm) (**Fig 2c**, solid line). This paradoxical response reflects a significant number of turns away from the wall, possibly triggered by the motion stimulus’ reflection off the acrylic surface of the wall whose mirrored direction contains motion energy in the opposite direction, when the wall is on the side of the fish.

To isolate sensorimotor transformations from contextual influences, we designed stimuli where visual motion was restricted to a constrained area (∼3 cm radius) centered around the fish (**Fig 2b** right**, S2 Movie**). Both species showed significantly improved turning responses to such restricted sine-grating stimuli compared to whole-arena stimuli (**Fig 2c** bottom**, d** dotted line). In zebrafish, this restricted stimulus effectively eliminated the wall effect (**Fig 2d** blue dotted line**, e** bottom, blue). However, in medaka, the wall effect persisted (orange arrowhead in **Fig 2d**), as demonstrated by continued position-dependent turning behavior even against a black background where the dark bottom and walls appear as continuous darkness. These results suggest that zebrafish are less influenced by peripheral visual information compared to medaka.

To systematically examine how the two species respond to motion at different spatial scales, we measured responses to dot motion stimuli covering varying spatial extents (**Fig 2f**). To that end we projected stationary flickering dots across the entire arena, where only a subset of dots within a restricted circle moved coherently relative to the fish. This arrangement allowed us to systematically vary the size of the motion area and thereby the optical angle subtended by the motion stimulus. Analysis of response magnitude versus motion area revealed distinct spatial integration profiles (**Fig 2g, h**). Zebrafish responded robustly to moving dot stimuli even at small optical angles (20 degrees), with response rates saturating at 99% already at 40 degrees. In contrast, medaka larvae exhibited a more gradual increase in turn angles as the spatial extent of moving dots expanded. Even at 80 degrees subtended visual angles medaka only reached an estimated saturation of 95% of turning, indicating that the effective visual angle might extend to regions above the horizontal plane (> 90 degrees).

To quantify these species differences, we calculated the motion energy of individual dots within the stimuli and integrated this energy over the motion area. We then fitted the relationship between integrated motion energy and larval turning responses to determine the spatial extent over which each species integrates motion information (**Fig 2i, S2 Fig**). By measuring the 95% threshold required to reach saturation response levels, we found that medaka demonstrated significantly broader spatial integration (threshold medaka tM = 32.8 ± 12.9 mm compared to zebrafish (threshold zebrafish tZ =17.8 ± 4.8 mm, (Mean ± STD)) (**Fig 2j, S2 Fig**). This quantitative analysis confirms that medaka require motion signals from a larger spatial area to achieve equivalent behavioral responses, reflecting fundamental differences in visual-motor integration strategies between the two species.

As a final validation of species differences in spatial integration, we presented spatially conflicting whole-field motion stimuli (**Fig 2k–n**). To that end sine-grating stimuli moving in one direction were projected beneath the fish, while opposing-direction gratings were projected in the surrounding area (**Fig 2k**). By varying the inner disk size, we determined the spatial extent required for local motion to balance global motion signals. When the inner disk was small, larvae turned in the direction of the outer (global) motion. As the inner disk size increased, larvae switched to following the inner (local) motion direction. The critical transition occurred at 23 degrees optical angle in zebrafish but at 39 degrees in medaka (**Fig 2m, n**), confirming that medaka require a larger proximal motion area to override global motion signals.

Together, these data demonstrate that medaka have broader spatial integration compared to zebrafish, consistent with a significantly wider field of view for motion processing.

### Temporal integration of flickering dot motion stimuli

Next, to investigate species differences in temporal motion integration, we employed random flickering dot stimuli and systematically varied stimulus lifetimes (**Fig 3**). We presented 500 small dots beneath freely swimming larvae, with dots moving either leftward or rightward according to the fish’s head orientation. Each dot had a limited lifetime and appeared at random positions within the restricted area beneath the fish (**Fig 3a**). Consistent with previous findings [14], zebrafish responded robustly to dot motion stimuli even with very short lifetimes (100 ms) (**Fig 3b, c**, blue**, S3 Movie**). In contrast, medaka exhibited significantly smaller turning responses to short-lifetime dot motion stimuli (**Fig 3b, c**, orange). However, as dot lifetime increased, medaka correct response ratios progressively improved to levels comparable to zebrafish. To quantify these temporal dynamics, we fitted exponential growth functions to the relationship between dot lifetime and response magnitude. This analysis revealed significant differences in temporal integration constants between species (**Fig 3d**). Zebrafish demonstrated rapid response saturation with integration times as short as 0.091 ± 0.013 (Mean ± STD) seconds, while medaka required longer stimulus durations (2.26 ± 0.23 seconds) to achieve equivalent response levels.

**Figure 3.**
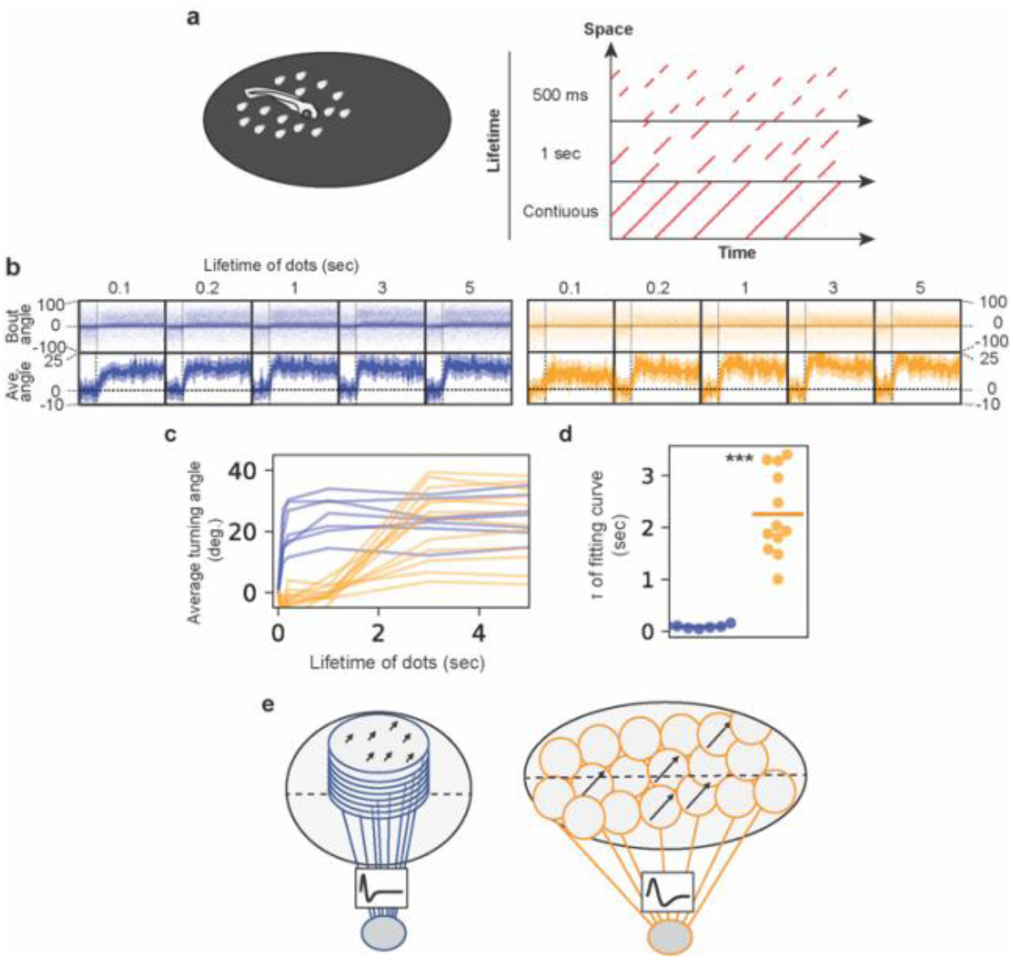
Comparative study in temporal domain in OMR using flickering dot motion. (a) Flickering dot motion stimuli presented to free-swimming larvae in a restricted area beneath the fish (left). Dots have defined lifetimes and random spatial locations (spatiotemporal patterns shown on right). (b) Turn responses to varying dot lifetimes during trials. Raw bout data (top) and averaged turn angles (bottom). Blue: zebrafish (n=7), orange: medaka (n=12). (c) Average turn angle increases more slowly in medaka (orange) than zebrafish (blue). Solid lines: experimental data. (d) The time constant of the fitting curve was shown. The values of each fish are shown dots and the averages are shown in horizontal lines. The time constants were significantly larger in medaka than zebrafish (zebrafish: 0.09 ± 0.01 seconds, medaka: 2.26 ± 0.23 seconds (Mean ± STD), Welch’s t-test, p=0.000001). (e) Schematic model of how visual stimuli could be processed in zebrafish (left) and medaka (right) larvae. In zebrafish, we hypothesized that the motion stimuli is averaged globally in a big receptive field (blue circle) and the top part of the retina. On the other hand, in medaka, we hypothesized that the motion stimuli is processed in a local receptive field which is tiled up in a bigger retina and temporally filtered and passed to the averaging step.

Based on these temporal integration profiles, we propose distinct underlying neural architectures. Zebrafish appear capable of rapidly integrating short-lifetime motion stimuli, potentially through retinal ganglion cells or downstream visual areas with larger receptive fields that enable efficient temporal summation of whole field motion stimuli. In contrast, medaka’s requirement for longer stimulus durations to achieve strong responses suggests a visual system composed of retinal ganglion cells with smaller receptive fields that integrate motion stimuli through longer temporal filters (**Fig 3e**).

### Divergent temporal filters uncovered by pulsed motion stimuli

To quantitatively estimate temporal filter properties for motion processing, we presented unidirectional motion pulses to fish larvae of both species and analyzed the dynamics of turning responses during and after the motion pulse (**Fig 4**). Using pulse stimuli of varying durations (**Fig 4a, b, S3 Fig**), we characterized both response onset and decay kinetics.

**Figure 4.**
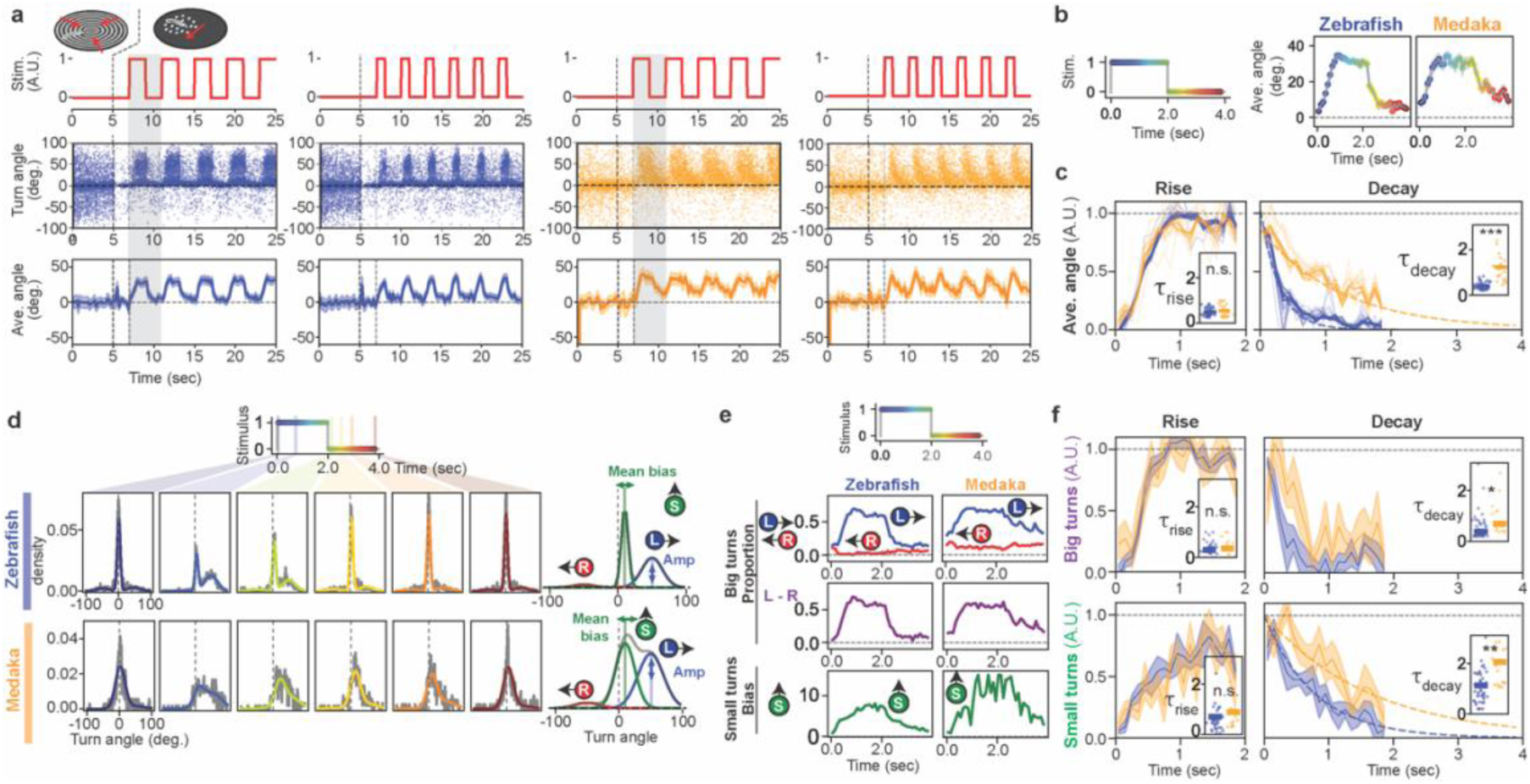
Divergent responses to unidirectional motion pulse stimuli reveal species-specific temporal dynamics. (a) Turn dynamics to unidirectional pulse stimuli. Trials begin with 5 s converging stimuli (black dotted lines), followed by dot-motion pulse stimuli (red trace, top). Individual (middle) and averaged (bottom) turn angles for zebrafish (blue, n=52) and medaka (orange, n=21; shaded areas: standard error). (b) Single-cycle stimulus-response analysis. One complete cycle is shown for zebrafish (left) and medaka (right). Dot colors correspond to temporal bins. (c) Normalized pulse responses and kinetics. Time zero in the ‘Rise’ plot indicates when the motion started, while time zero in the ‘Decay’ indicates when the motion stopped. Rise time constants: no species difference (zebrafish: 0.48 ± 0.02 seconds, medaka: 0.55 ± 0.06 seconds; p=0.52, n.s., mean ± SEM; Mann-Whitney U test). Decay time constants: significantly longer in medaka (zebrafish: 0.41 ± 0.02 seconds, medaka: 1.24 ± 0.14 seconds, mean ± SEM; p=4.0×10⁻¹⁰, Mann-Whitney U test). Blue: zebrafish, orange: medaka. Thin lines represent temporal dynamics of normalized average turn angles for individual fish; thick lines represent population averages. Inset dot plots show individual fish time constants for rise and decay phases. (d) Three-component Gaussian decomposition: large leftward (blue), large rightward (red), and small turns (green). Temporal dynamics in 100 ms bins with fitted curves overlaid. (e) Temporal evolution of behavioral components. Large-turn amplitudes (top), small-turn means (bottom), and large-turn asymmetry (left minus right amplitude, middle, purple). (f) Species comparison of temporal dynamics. Time constants (mean ± SEM): zebrafish big turns (rise: 0.302 ± 0.004 seconds, decay: 0.384 ± 0.037 seconds), zebrafish small turns (rise: 0.701 ± 0.04 seconds, decay: 1.130 ± 0.04 seconds, medaka big turns (rise: 0.363 ± 0.014 seconds, decay: 0.684 ± 0.141 seconds), medaka small turns (rise: 0.894 ± 0.096 seconds, decay: 2.041 ± 0.059 seconds). Zebrafish n=53, medaka n=20. Statistical comparisons (Mann-Whitney U test): big turn rise p=0.93, big turn decay *p=0.013, small turn rise p=0.13, small turn decay **p=0.0067 (asterisks denote *p<0.05, **p<0.01).

Analysis of average turning responses revealed similar onset time constants across species (zebrafish: 0.48 ± 0.02 seconds, medaka: 0.55 ± 0.06 seconds), but significantly longer decay time constants in medaka (1.24 ± 0.14 seconds) compared to zebrafish (0.41 ± 0.02 seconds) (**Fig 4c**). This difference suggests that while both species initiate responses with comparable speed, medaka maintain motion-driven turning behavior for substantially longer periods after stimulus offset.

Previous studies show that zebrafish larvae exhibit two distinct types of turning behavior—small turns and big turns—which are readily distinguishable in turn angle distributions [9]. In medaka, the distributions are less obviously fit with gaussian distributions compared to zebrafish, but still we can fit with three gaussian distributions and perform similar analysis (see also **S4 Fig**). To investigate the temporal dynamics of these behavioral components separately, we generated turn angle distributions for each 100 ms time bin during stimulus presentations (**Fig 4d, 4e**) and fitted them with three-component Gaussian distributions. We extracted the difference between big left and big right turn frequencies as the “big turn proportion” (**Fig 4e** middle, purple) and the mean bias angle of small turns as the “small turn bias” (**Fig 4e** bottom, green). Exponential curve fitting of both “big turn proportion” and “small turn bias” revealed that rise and decay time constants of big turns were not significantly different between zebrafish and medaka (**Fig 4f**, top left). This suggests that the neural circuits processing signals inducing large-amplitude turning maneuvers have similar temporal characteristics across species, potentially reflecting conserved mechanisms for rapid escape or orientation responses.

In contrast, small turns displayed markedly different temporal profiles. Both species showed longer rise time constants for small turns (zebrafish 0.701 ± 0.04 seconds, medaka 0.894 ± 0.096 seconds) compared to big turns (zebrafish: 0.302 ± 0.004 seconds, medaka: 0.363 ± 0.014 seconds) (**Fig 4f**, left column), indicating low pass filtering of signals inducing small turns. Most strikingly, decay time constants for small turns exceeded rise time constants in both species, but medaka exhibited significantly longer small turn decay constants compared to zebrafish (zebrafish: 1.130 ± 0.04 seconds, medaka: 2.041 ± 0.059 seconds) (**Fig 4f**). This represents a remarkable species difference, with medaka small turn responses persisting nearly twice as long as those in zebrafish. Given that neural spiking occurs on millisecond timescales, this ∼1-second difference in behavioral persistence suggests fundamentally different temporal integration mechanisms underlying small turn control between the two species.

Since small turns play a more dominant role in medaka swim behavior than in zebrafish, this extended time constant becomes prominent also in the overall average turn angle (**Fig 4c**), which relaxes in medaka with a time constant that is three times longer than that of zebrafish (medaka: 1.24 ± 0.14 seconds, zebrafish: 0.41 ± 0.02 seconds).

### Temporal integration of alternating stimuli

The prolonged decay kinetics observed in medaka’s average turning responses and small turn components suggested enhanced signal retention capacity compared to zebrafish. To test this hypothesis systematically, we presented motion stimuli that alternate leftward and rightward across a range of frequencies and performed frequency-domain analysis.

Even in zebrafish, which showed relatively short decay time constants, the response curves deviated from pure sinusoidal patterns, exhibiting steep slopes that indicated nonlinearities in the underlying control systems (**Fig 5a, b**). Frequency-amplitude analysis revealed distinct processing strategies between species: zebrafish demonstrated approximately linear amplitude decay with increasing stimulus frequency, while medaka exhibited exponential amplitude decay (**Fig 5c**). This difference suggests fundamentally different frequency-dependent filtering mechanisms.

**Figure 5.**
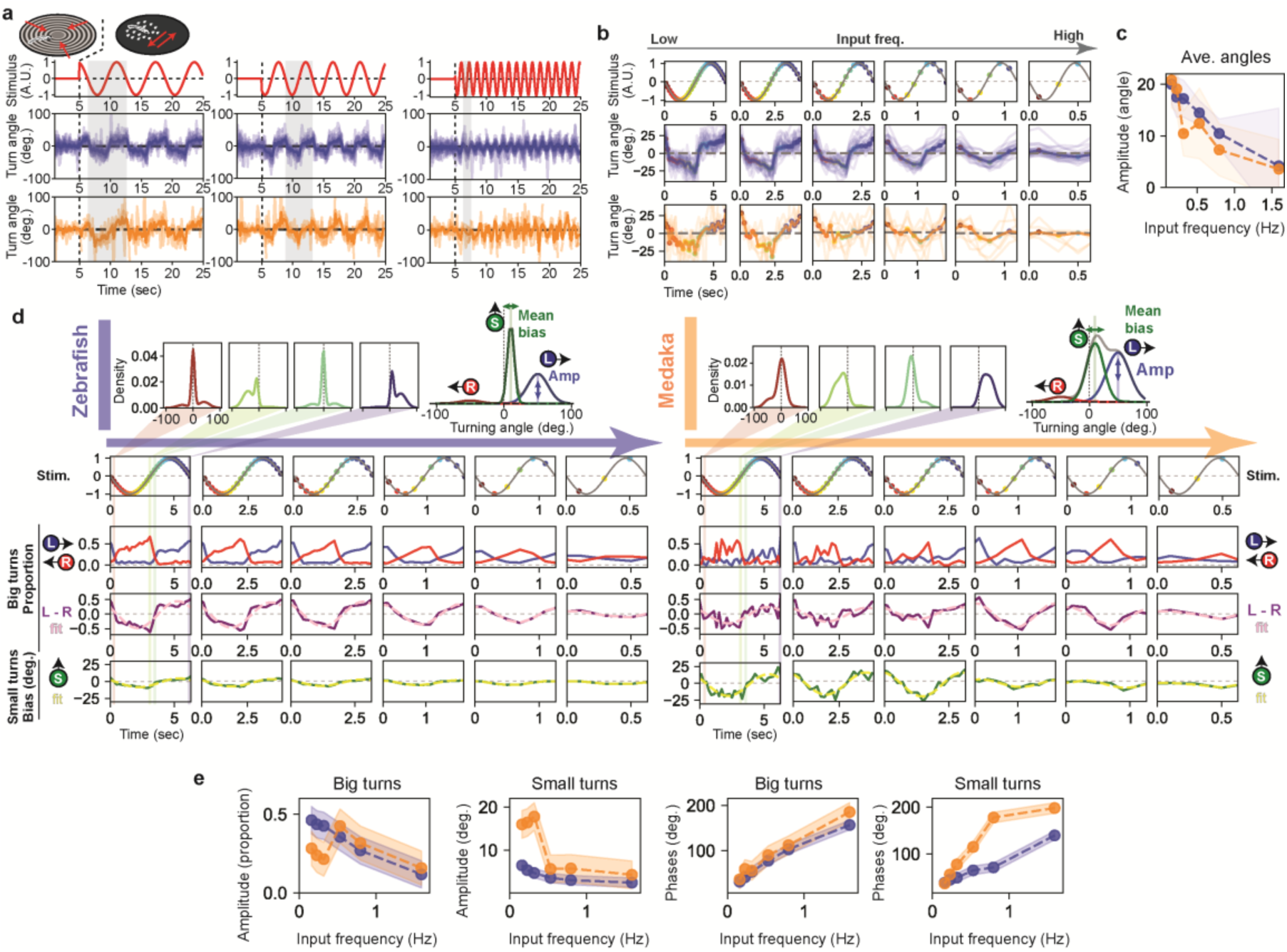
Optomotor response to the alternating stimuli. (a) Turn dynamics to alternating stimuli. Each trial begins with converging stimuli for 5 seconds (indicated by black dotted lines), followed by presentation of moving dots that alternate between leftward and rightward motion with velocities following a sinusoidal pattern (red, top row). (b) Frequency-response characterization. Average turn angles plotted with colored dots indicating time bins within cycles and corresponding responses. Input and response frequencies show a linear relationship. (c) Response amplitude analysis. Amplitudes of fitted sine waves plotted versus stimulus frequency. (d) Turn angle distribution analysis within cycles. Fitted curves for 200 ms bins during one complete cycle (2π) for zebrafish (left) and medaka (right). Line colors correspond to temporal positions. Three Gaussian components are fitted: large rightward (red), large leftward (blue), and small turns (green) (top right). Amplitudes of large-turn distributions (third row) and means of small-turn distributions (bottom, green) across cycles. Large-turn asymmetry dynamics (left minus right amplitude, fourth row, purple) and small turns fitted with sine waves (dotted pink: large turns, dotted yellow: small turns). (e) Frequency-domain analysis of behavioral components. Following sine wave fitting, amplitude and phase of large and small turn components plotted to characterize frequency-amplitude and frequency-phase response profiles. Shaded regions: confidence intervals (blue: zebrafish, orange: medaka).

To dissect the temporal dynamics of big turns and small turns separately, we fitted three-component Gaussian distributions to turn angle distributions within each 200 ms time bin (**Fig 5d, S4 Fig**). Frequency-amplitude analysis of these components revealed species-specific patterns. Zebrafish showed relatively linear amplitude decay across frequencies for both big turns and small turns (**Fig 5e**, left blue), indicating uniform frequency processing across behavioral components. In contrast, medaka exhibited more complex frequency dependencies. Most notably, medaka big turns showed amplitude peaks at lower stimulus frequencies (∼0.5 Hz) (**Fig 5e**, left orange), suggesting resonance or optimal frequency tuning in their large-amplitude turning system.

Frequency-phase analysis provided additional insights into temporal processing mechanisms. For big turns, zebrafish and medaka showed no significant phase differences across frequencies, consistent with the similar time constants observed for big turn responses to unidirectional pulse stimuli. However, small turn phase analysis revealed substantial differences between species (**Fig 5e**, right), with medaka showing significantly larger phase lags that increased with frequency. This pattern is consistent with the much longer decay time constants observed for medaka small turns in the pulse stimulus experiments, confirming that species differences in temporal integration are primarily mediated through small turn control mechanisms rather than big turn systems.

These frequency-domain results support a model where medaka have evolved slower temporal integration for fine motor adjustments (small turns) while maintaining rapid response capabilities for large corrective maneuvers (big turns), potentially reflecting adaptations to their clearer, more spatially complex visual environments.

### Mechanistic models reproduce species-specific temporal dynamics

To test our understanding of the underlying neural mechanisms, we developed a mechanistic computational model that simulates the temporal dynamics of optomotor responses using the Euler method, and compared the simulated results with actual data (**Fig 6a**). The model implements separate control systems for big turns (proportional control) and small turns (bias control), each governed by input gain (g₀) and decay rate (g₁) parameters. Based on observations from alternating stimuli experiments, we also incorporated a softmax function to capture the nonlinear response characteristics observed in both species.

**Figure 6.**
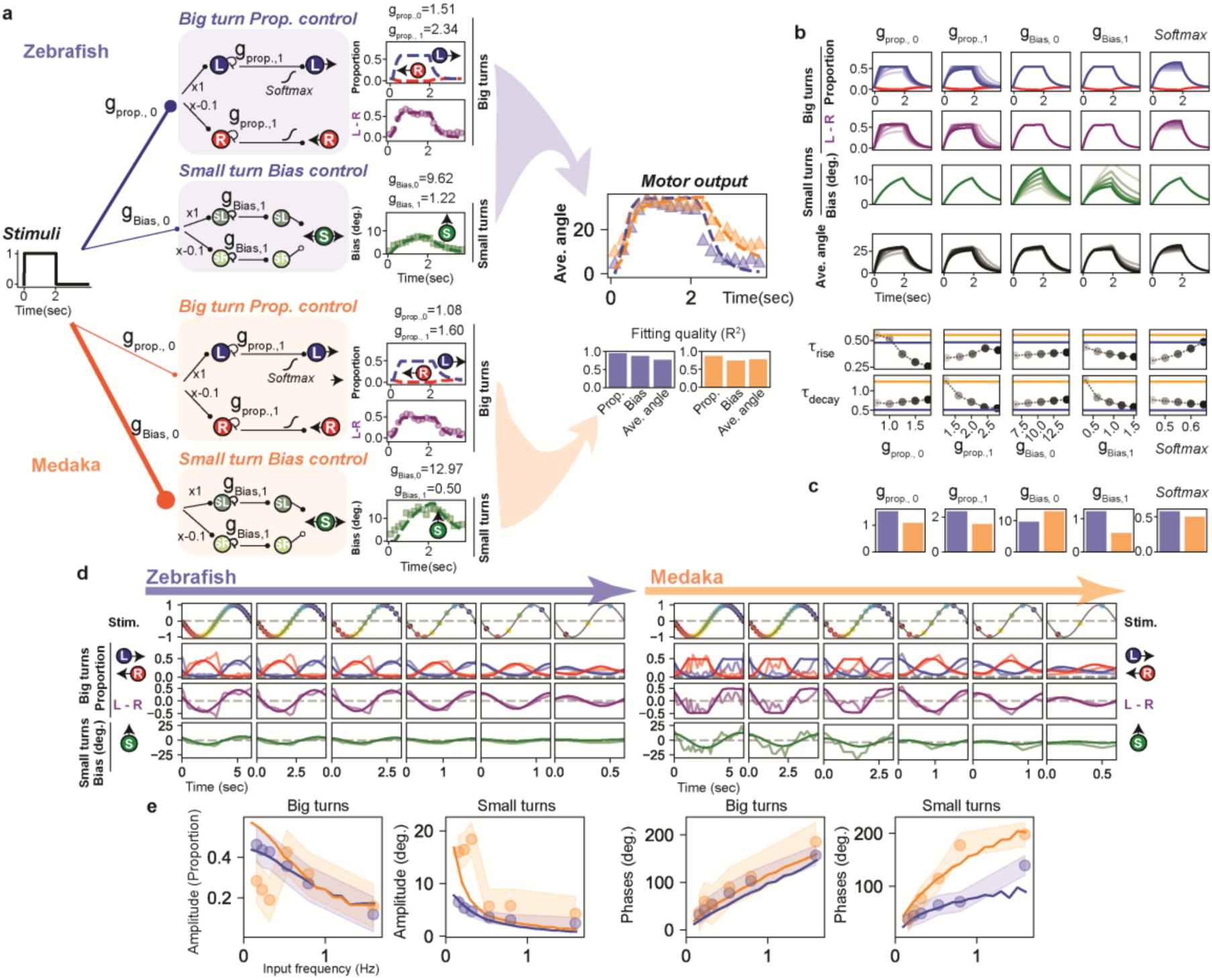
Temporal integration model captures species-specific optomotor responses. (a) Model architecture and fitting. Schematic shows sensorimotor control for zebrafish (blue) and medaka (orange). Visual input drives two pathways: proportion control (big turn probabilities) and bias control (small turn direction). Each implements first-order dynamics with gain (g0) and decay (g1), followed by saturation. Components shown: big turns left (blue), right (red), difference (purple), and small turn bias (green). Fitted parameters displayed above plots. Dashed lines: model predictions; dots: experimental data. Average angles (right) emerge from component combinations. Bar graphs show R² fitting quality. (b) Parameter sensitivity analysis. Model responses across parameter ranges (proportion gain g₀, decay g₁, bias gain fg₀, decay fg₁, saturation). Top: temporal dynamics with line shading corresponding to parameter values (x-axes, bottom). Bottom: rise and decay time constants versus parameters. Horizontal lines: experimental values (blue: zebrafish, orange: medaka). (c) Species-specific parameter estimates. Fitted values reveal distinct temporal processing strategies between species. (d) Simulated dynamics for alternating stimuli using parameters from (c). Experimental data from Figure 5 (shaded lines) compared to simulations (darker lines). (e) Frequency domain validation. Model predictions (dashed) using parameters from (c). Amplitude and phase responses versus frequency for big turn difference and small turn components. Experimental data (shaded: mean ± CI) validate predictions, revealing species differences: zebrafish show linear amplitude decay while medaka exhibits sharp cutoff around 0.5 Hz.

Analysis of the three-component Gaussian distributions revealed that medaka small turns contribute substantially more to average turning behavior than zebrafish small turns, a feature we incorporated into our modeling framework. Parameter sensitivity analysis demonstrated how variations in big turn proportions (purple) and small turn bias (green) influence overall behavioral outputs across different parameter ranges (**Fig 6b**).

Fitting the model to experimental data from both species revealed key parameter differences (**Fig 6c**). Most notably, decay rates (g₁) were consistently lower in medaka compared to zebrafish for both big turn and small turn control systems. This finding directly corresponds to the longer temporal integration observed in medaka across multiple experimental paradigms, providing mechanistic support for species differences in motion processing persistence.

To test the model, we fitted parameters based on the unidirectional stimulus responses (**Fig 4**) and tested the model by predicting responses to alternating stimuli (**Fig 5d, 6d**). Frequency domain analysis applied to these predictions confirmed most key experimental features **(Fig 5e, 6e)**. Specifically we could capture the steep exponential decay in small turn frequency-amplitude responses observed in medaka, and a more linear decay in those of zebrafish. We also captured the progressive phase delay with increasing input frequencies in both turn modes in zebrafish. However, the model failed to capture the transient increase in big turns at intermediate frequencies in medaka. In addition, the persistent high amplitude in small turns for the lower frequencies is not captured by the model who predicts a consistent and exponential decrease in small turn bias with increasing stimulus frequencies.

These modeling limitations suggest the presence of more complex, frequency-dependent mechanisms that are not captured by simple first-order linear dynamics. Specifically, the discrepancies point toward potential behavioral switching between big turn and small turn control systems depending on input frequency, with medaka possibly employing different processing strategies at low versus high frequencies. This frequency-dependent mode switching could represent an adaptive mechanism for optimizing responses to different timescales of environmental motion, warranting future investigation of more sophisticated control architectures.

## Discussion

In this study, we compared how zebrafish and medaka larvae integrate whole-field motion across space and time. Despite their similar morphologies and behavioral repertoires, the two species exhibited pronounced differences in both spatial pooling and temporal retention. Medaka integrated motion over larger visual fields and showed stronger sensitivity to peripheral cues, whereas zebrafish relied more heavily on motion directly beneath the body. Temporally, both species shared similar response onsets, but medaka exhibited markedly longer persistence of motion-driven activity, suggesting enhanced temporal retention at the cost of responsiveness to rapidly fluctuating stimuli.

### Spatial Integration Differences

Our spatial analyses indicate that zebrafish primarily extract motion below their bodies, whereas medaka combine motion signals across a broader spatial domain, potentially extending above the horizontal plane. This difference was especially evident in wall-proximity assays, where medaka exhibited substantial decrements in turning performance near boundaries, which can be caused by allocation of visual attention across vertical axes [25]. Such species differences may arise from variation in eye position, retinal topology, or receptive-field organization in early visual pathways. Indeed, zebrafish and medaka differ in photoreceptor distributions and mosaic patterns [26, 27], which could shape the effective spatial kernels driving OMR behavior.

### Temporal Integration and Object Detection Strategies

Flickering-dot experiments revealed strong divergence in temporal integration rules. Zebrafish reached peak performance at very short lifetimes (≤200 ms), whereas medaka required much longer durations (>2 seconds) for optimal responses. These results imply species-specific implementation of temporal filters, potentially through differences in retinal ganglion cell dynamics or downstream temporal integration mechanisms [28]. Medaka’s strong improvement with prolonged lifetimes aligns with enhanced sensitivity to persistent motion, a computation relevant for object permanence detection and sustained tracking of conspecifics [29, 30]. Conversely, zebrafish appear optimized for rapid extraction of transient motion cues, enabling quick orientation adjustments in fluctuating environments.

### Component-specific Temporal Dynamics

Pulse-response decompositions revealed asymmetric temporal dynamics specifically in medaka which showed significantly slower decay than rise times. Importantly, small turns exhibited significantly longer time constants than large turns in both species, indicating partially dissociable control modules. Medaka showed especially prolonged persistence in small-turn components (∼2 seconds vs. ∼1 seconds in zebrafish), suggesting species differences in circuits governing fine motor adjustments. This aligns with known reticulospinal architecture in zebrafish where nucleus of the medial longitudinal fasciculus (nMLF) neurons and associated hindbrain circuits selectively regulate small corrective turns [31–33]. Comparative physiological and anatomical studies of these circuit nodes could help reveal how temporal retention mechanisms diverge across species.

### Evolutionary and Social Implications

The distinct temporal and spatial integration profiles observed here likely reflect evolutionary tuning to different ecological and social demands. Medaka, which show improved social approach in juveniles [30] and exhibit stronger schooling tendencies in adults than zebrafish [34], may benefit from enhanced temporal persistence and broader spatial pooling to maintain stable trajectories and monitor neighbors during coordinated group motion [35]. Zebrafish, which form less aligned shoals, may rely on rapid, transient integration to support agile maneuvering and quick adaptation to unpredictable flow conditions. Although the ancestral environments of these species cannot be reconstructed directly, differences in morphologies of the chorion, reproductive habits [36, 37], early locomotor mobility [38], and social responsiveness support the broader view that distinct ecological and social pressures shaped their motion-processing algorithms.

### Mechanistic Modeling and Computational Principles

Our computational model captured major species differences using minimal changes in gain and decay parameters, suggesting that a small number of circuit-level motifs, such as altered recurrent connectivity [39–41] or neuro-modulatory tone [42], can account for divergent behavioral dynamics. The model’s inability to fully reproduce medaka’s low-frequency response patterns points toward additional nonlinear or frequency-dependent mechanisms [43], possibly reflecting behavioral switching or higher-order control loops [44]. These predictions can be directly tested in future electrophysiological, imaging, or connectome studies [45].

Together, our findings reveal that zebrafish and medaka rely on distinct computational strategies for processing visual motion, emerging from differences in spatial pooling, temporal retention, and component-specific integration dynamics. Despite outwardly similar behaviors, the underlying algorithms diverge substantially, highlighting how evolution can reshape core sensorimotor computations even within related vertebrate lineages.

## Materials and Methods

### Ethics statement

All experiments followed institution IACUC protocols as determined by the Harvard University Faculty of Arts and Sciences standing committee on the use of animals in research and teaching. The animal experimentation protocols, 25-03 and 22-04 were submitted and approved by this institution’s animal care and use committee (IACUC). This institution has Animal Welfare Assurances on file with the Office for Laboratory Animal Welfare (OLAW), D16-00358 (A3593-01).

We followed the ARRIVE guideline 2.0; the sample size is described in the method section. Inclusion and exclusion criteria are also mentioned in the method section. For each experiment, different visual stimuli were shown in random order. Data was analyzed by blinded and masked.

### Fish maintenance

Wild-type zebrafish (WIK strain) and medaka (dr-R strain) were used in this study. Embryos were incubated at 27 °C. Zebrafish hatched within 2–3 days, while medaka hatched after 7–8 days. Prior to hatching, embryos of both species were maintained in embryo water supplemented with methylene blue. After hatching, larvae were reared exclusively in filtered fish facility water (200 nm pore size), with water changes every other day. Based on published literature and the timing of locomotor development, experiments were conducted on zebrafish at 5–6 days post fertilization (dpf) and on medaka at 1 day post hatch (dph), unless otherwise specified.

### Free-swimming behavioral experiments

Free-swimming behavioral experiments were conducted on fish larvae following previously established protocols [14, 23, 25]. Briefly, behavior was recorded at 90 frames per second using an overhead camera (Grasshopper3, FLIR). Sandblasted dishes (radius: 6cm) were used to present visual stimuli to the larvae [46]. To extract behavioral features, the mode image of the first 1-minute segment of each recording was subtracted, allowing the center of mass and head angle of the focal fish to be tracked at 90 frames per second. Each experimental epoch began with a 5-second presentation of an open-loop converging stimulus, followed by 20 seconds of closed-loop visual stimulation. Visual stimuli were projected from below at 60 frames per second using an AAXA P300 projector. A single trial consisted of a randomized sequence of different visual stimuli, and each experiment included 15 repeated trials.

### Visual stimuli

Visual stimuli were projected from below the behavioral rig and programmed in Python using the Panda3D package. Each epoch began with a 5-second open-loop presentation of converging stimuli (speed: 0.15, wavelength: 0.15, contrast: 1). The luminance of the stimulus, represented as a grayscale value C, was defined by the equation (r: radial position in the disk, t: time, v: velocity, K: constant):

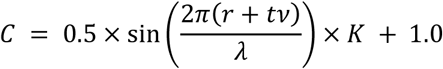

Following the open-loop phase, closed-loop stimuli were presented for 20 seconds. These included either sine-grating or random dot motion stimuli. For sine-grating stimuli (speed: 0.15, wavelength: 0.15, contrast: 1), the luminance C was defined as:

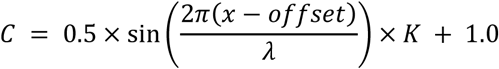

where x is the position of the fish. Random dot motion stimuli (Fig 3) were generated following previous work [14]. For restricted stimuli, the visual stimulus disk was reduced to half the radius (number of dots: 400). During closed-loop presentation, the stimulus disk rotated in real time based on the fish’s measured head angle. For whole-field stimuli (Figs 1 and 2), one of four motion directions (forward, leftward, rightward, or backward) was randomly selected and shown. In the restricted condition (Fig 2), the stimulus disk was scaled to half the rig size and centered on the fish’s center of mass. The area outside the disk was black. In the variable-disk-size dot motion stimuli (Fig 2f–j), dot density and size were kept constant. For the inconsistent stimuli (Fig 2k–n), the outer disk had a radius of three-quarters of the full field, and an inner disk displayed a sine-grating moving in the opposite direction. The size of the inner disk was randomly chosen, and each fish experienced 15 trials. In Fig 3, the lifetime of dots was randomly chosen from the set (100 ms, 500 ms, 1 second, 2 seconds, 4 seconds, 10 seconds), while keeping the total number of visible dots constant throughout the session. For the pulse stimuli (Fig 4) and alternating stimuli (Fig 5), dot stimuli had a fixed lifetime of 20 seconds. In pulse stimuli experiments, the 5-second converging stimulus was followed by a 2-second rest period, after which a series of pulse and rest periods was presented. Ten stimulus conditions (combinations of five pulse durations and two rest durations) were randomly ordered per trial. Each fish experienced 15 trials. In the alternating stimuli experiment (Fig 5), leftward and rightward dot motion pulses were shown alternately. The speed of dot motion followed a sine-wave profile.

### Data inclusion and exclusion criteria

For the free-swimming behavioral experiments shown in Fig 1 and Fig 2a, all fish were included in the analysis. However, we observed that fish swimming near the walls exhibited reduced responses to motion stimuli. Therefore, for the experiments shown in Figs 3–5, we excluded fish that tended to swim near the walls. Specifically, we calculated the average distance between each fish’s position and the center of the arena across the entire experiment. Fish with an average distance greater than 3.5 cm were excluded from the analysis.

### Calculating turning angle in larval teleost & ratio of correct turn

To quantify turning behavior in larval fish, we first measured the displacement of the center of mass over time to compute swimming speed. Peaks in the speed profile were detected and defined as “turn” events. For each detected turn, the change in head angle before and after the speed peak was calculated as the turn angle. Throughout the experiments, we recorded the turn angle, timing, and spatial position of each turn within the behavioral arena for further analysis. Turn angles were averaged across all turning events that occurred during sine-grating or dot motion stimuli, computed in 100 ms time bins. Both leftward and rightward motion stimuli were presented during the experiments. For presentation purposes, responses to rightward stimuli were mirrored and combined with responses to leftward stimuli to generate a unified representation. To calculate the correct turning ratio, a turn was classified as a “correct turn” when the sign of the fish’s turning angle matched the direction of the visual motion stimulus (leftward: positive; rightward: negative). The correct ratio was defined as the number of correct turns divided by the total number of turns. A correct ratio of 0.5 indicates random turning behavior.

### Calculation of motion energy in different size of motion disks and estimation of thresholds in teleost larvae

To estimate and compare motion energy across different sizes of dot-motion disks (Fig 2f–j), we first computed the retinal occupancy of each dot (radius: *r*) located at position (*x*, *y*) for a fish swimming at height *h*. Specifically, we calculated the horizontal (φ) and vertical (θ) angular spans subtended by each dot on the retina (see S2 Fig a, b). We then computed the rate of change of these angles (dφ/dt and dθ/dt) over time as the dots moved along the *x*-axis (S2 Fig b). The motion energy contributed by each dot in the *x–y* plane, denoted as *E_abs_(x, y)*, was defined as the product of dφ/dt and dθ/dt (S2 Fig c). The total motion energy within a given motion disk (radius: *R*), denoted as *E_total_(R)*, was calculated by summing the visual motion energy over the entire disk area (S2 Fig d). To estimate motion sensitivity thresholds in zebrafish and medaka larvae for different disk sizes, we fit the behavioral responses to an exponential curve. The threshold was defined as the radius at which the behavioral response reached 95% of its asymptotic (maximum) value.

### Estimation of time constants

To investigate species differences in temporal integration of motion stimuli (Fig 4), we presented unidirectional visual pulse motion stimuli to free-swimming zebrafish and medaka larvae and quantified their turning responses. Each trial began with 5 seconds of open-loop converging stimuli, followed by a randomly selected combination of motion and resting durations. This pulse pattern was repeated for a total of 25 seconds. Motion durations were 5, 2, 1, or 0.5 seconds, and resting durations were either 2 or 0.25 seconds. Each fish underwent 15 trials.

To analyze the temporal dynamics of the turning response, all turns from the repeated pulses were aggregated per fish and treated as a single composite sequence. The rise and decay phases of the averaged turn angle were then quantified by fitting exponential functions and extracting the time constant (τ) as follows:

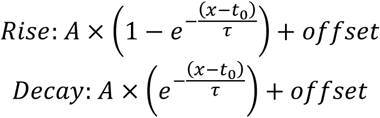

where *A* is the amplitude and *t₀* is the onset time.

For statistical analysis, normality of data distribution was assessed using the Shapiro–Wilk test, and homogeneity of variances using Levene’s test. Based on these results, the Mann–Whitney U test was used to compare between species. Effect sizes were calculated using Cohen’s *d*. Data are presented as mean ± standard error of the mean (SEM), and statistical significance was set at *p* < 0.05.

To further examine the temporal dynamics of large and small turns during pulse stimuli, turn angle distributions were generated in 100 ms bins for Fig 4 (and in 200 ms bins for Fig 5), and fitted using a mixture of three Gaussian distributions:

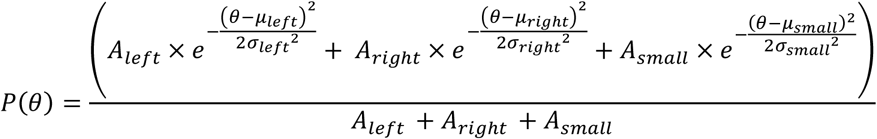

As turn characteristics differ by species, we used the following parameters for fitting: Zebrafish: μ_right_= –40.0, μ_left_= 40.0, σ_right_= 20.0, σ_left_= 20.0, σ_smal_= 6.0; Medaka: μ_right_= –45.0, μ_left_= 45.0, σ_right_= 20.0, σ_left_= 20.0, σ_smal_= 15.0. From the fits, we extracted the amplitude of large turns (Amp_left_, Amp_right_) and the directional bias of small turns (μ_small_). The temporal dynamics of these parameters were then analyzed by fitting exponential curves to estimate time constants.

### Alternative stimuli and frequency domain analysis

Sinusoidal motion stimuli were generated with dot velocities modulated according to *v(t)* = *v*₀ × sin(2π*ft*), where *v*₀ = 10 mm/s maximum velocity amplitude and *f* was the stimulus frequency (positive: leftward, negative: rightward). Stimulus frequencies were tested at (1/0.1)*2π, (1/0.2)*2π, (1/0.3)*2π, (1/0.5)*2π, (1/0.7)*2π, and 2π Hz. Each frequency condition was presented for 20 seconds after 5 seconds of open-loop converging stimuli. Stimulus order was randomized across trials. Turning angles were calculated as described in the “Behavioral Analysis” section. For each frequency condition, mean turning angles were computed across all individuals within a 200ms time bin. Then time series of average turning angles were fitted with the function *θ(t)* = *A* × sin(2π*ft* + *φ*) + *θ*₀ using least-squares optimization (scipy.optimize.curve_fit). Response amplitude (*A*) and phase lag (*φ*) were extracted for each frequency condition. To analyze the temporal dynamics of big turns and small turns, for each 200 ms time bin, individual fish turn angle distributions were generated and fitted with three-component Gaussian mixture models as described in the “Estimation of time constants” section. This yielded temporal traces of: big turn proportion (L-R) and Bias of small turn. Temporal dynamics of big turn proportions and small turn bias were fitted with sinusoidal functions using the same procedure as for average turning angles. Component-specific amplitude and phase responses were extracted to characterize frequency dependencies of underlying control systems.

### Modeling and simulation

To simulate the temporal dynamics of optomotor responses to whole-field motion, we developed a mechanistic model based on hypothesized neural control circuits in fish larvae. The model incorporates two independent control systems: (1) proportional control, governing big turn probabilities and (2) bias control, governing small turn direction.

Each control system implements first-order dynamics with two key parameters: input gain (*g*₀) determining sensitivity to visual stimuli, and decay rate (*g*₁) controlling the temporal persistence of neural activity. The temporal evolution of neural responses was computed using the Euler method with time step *dt* = 0.01 s:

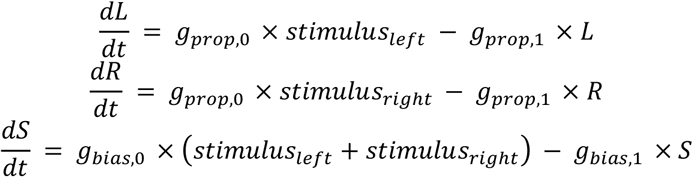

where *R* and *L* represent right and left big turn activities, and *S* represents small turn bias. Visual stimuli were processed asymmetrically: for leftward motion, *stimulus_left* = 1.0 and *stimulus_right* = –0.1; for rightward motion, these values were reversed.

In addition, we implemented transport delays (100 ms) for big turns in both species, which could arise from neural transport delays, second-order membrane dynamics, or a combination of both mechanisms.

Then, neural activities were converted to behavioral outputs through a three-component Gaussian mixture model as described in the “Estimation of time constants” section. Also, as a realistic model, we estimated that both left, right, and small turns control have a baseline activity. A saturation boundary was applied to prevent unrealistic neural activities, described as the “softmax” parameter in Fig 6.

### Parameter Fitting and Validation

Model parameters were fitted to experimental results of unidirectional stimuli using the component-based approach: we directly fitted the left-right difference (LRs = L – R) and small turn bias (S) components, allowing average turning angles to emerge naturally from the angle distribution function. This mechanistically sound approach ensures that fitted components correspond to hypothesized neural pathways.

Non-linear least squares optimization (scipy.optimize.minimize of python package) was used with the “L-BFGS-B” algorithm and multiple random initial conditions to avoid local minima. Fitting quality was assessed using the coefficient of determination (R²): R² = 1 – (SS_res / SS_tot) where SS_res is the sum of squared residuals and SS_tot is the total sum of squares. Separate R² values were calculated for fitted components (LRs, F) and emergent average angles.

To validate model predictions across different stimulus conditions, we tested the fitted parameters against alternating sinusoidal motion stimuli at frequencies ranging from 0.1 to 10 Hz. Model responses were analyzed in the frequency domain using Fourier analysis to extract amplitude and phase characteristics, enabling comparison with experimental frequency response data.

### Statistics

**(*Fig 2d,e*)** Turn angle data were analyzed using a two-way mixed analysis of variance (ANOVA) with visual stimuli (whole-rig stimuli vs. restricted stimuli) as a between-subjects factor and spatial bin (bins ‘0-‘ – ‘50-‘, from wall to center) as a within-subjects factor. Average turning angles calculated for each fish in each spatial bin. Following significant main effects or interactions, post-hoc analyses were conducted. For the main effect of spatial bin, pairwise comparisons with Bonferroni correction were performed to determine differences in turning angles between spatial locations. For the significant stimuli × spatial bin interaction, simple main effect analyses were conducted by comparing stimuli differences at each spatial bin using independent samples t-tests with Bonferroni correction for multiple comparisons. Effect sizes were reported as partial eta-squared (ηp²) for ANOVA effects and Cohen’s d for pairwise comparisons. Statistical significance was set at p < 0.05. All statistical analyses were performed using Python (version 3.10.13) with the pingouin (version 0.5.5) and scipy (version 1.10.1) packages.

**(*Fig 2j*)** Statistical analyses were performed using Python (version 3.10.13) with SciPy (version 1.10.1) and NumPy (version 3.10) libraries. Data are presented as mean ± standard deviation (STD). Sample sizes were n=10 for zebrafish and n=12 for medaka. The normality of data distribution in each group was assessed using the Shapiro-Wilk test. As both groups showed normal distribution (Shapiro-Wilk test, p > 0.05, zebrafish: 0.5894, medaka: 0.2263), statistical comparisons between groups were performed using Welch’s t-test, which does not assume equal variances between groups.

**(*Fig 2n*)** The cancel effect size was quantified as the zero-crossing point of the interpolated function of the inside-disk angle and turn angle. To compare the cancel effect size between zebrafish (n=6) and medaka (n=10), we calculated the average effect across 15 trials for each individual fish. The normality of data distribution was assessed using Shapiro-Wilk tests, and homogeneity of variances was evaluated using Levene’s test. Based on these results, Student’s t-test was used to compare between species. Effect size was calculated as Cohen’s d. Data are presented as the mean ± standard error of the mean (SEM). Statistical significance was set at p < 0.05. All statistical analyses were performed using Python Python (version 3.10.13), with scipy (version 1.10.1), statsmodels (version 0.14.0), and (version 0.5.5) packages.

**(*Fig 3*)** To compare the ability to respond to short lifetime of dots between zebrafish (n=7) and medaka (n=12), we calculated the average of the time constants across 20 trials for each individual fish. The normality of data distribution was assessed using Shapiro-Wilk tests, and homogeneity of variances was evaluated using Levene’s test. Based on these results, Welch’s t-test was used to compare between species. Statistical significance was set at p < 0.05.

**(*Fig 4*)** For comparisons between zebrafish and medaka, we used non-parametric Mann-Whitney U tests (two-tailed) due to non-normal distributions of behavioral parameters and unequal sample sizes between species. No corrections for multiple comparisons were applied, as each statistical test addressed a distinct hypothesis regarding species differences in temporal dynamics. Statistical significance was defined as *p < 0.05, **p < 0.01, ***p<0.001.

(***Fig 5***) Confidence intervals for fitted sine wave parameters were calculated using the covariance matrix obtained from the nonlinear least-squares fitting procedure. Parameter standard errors were derived as the square root of the diagonal elements of the covariance matrix. Confidence intervals were then computed using the t-distribution with degrees of freedom equal to the number of data points minus the number of fitted parameters (n – 3). The confidence interval half-width for each parameter was calculated as the product of the critical t-value (α = 0.05 for 95% confidence intervals) and the corresponding parameter standard error. For phase measurements, confidence intervals exceeding 360° were capped at this maximum value to reflect the periodic nature of the parameter. Fitting quality was assessed using the coefficient of determination (R²) and reduced chi-squared statistics, with the latter calculated as the sum of squared residuals divided by the degrees of freedom.

## Data access

Behavioral data is submitted to Dryad (DOI: 10.5061/dryad.gxd2547wg). Code for analysis is shared in the github link (https://github.com/yasukoisoe/cross_species_OMR).

## Acknowledgments

We are grateful to the Engert lab and the Fishman lab at Harvard University for experimental and conceptual support. We thank Dr. Roy Harpaz at Harvard University for critical discussion. We thank Dr. Armin Bahl at Konstanz Institute for experimental support and thoughtful suggestions in behavioral rigs and analysis.

## Author Contributions

Conceptualization, YI, and FE; Methodology, YI, FE; Data analysis, YI; Statistics, YI, YM, MB; Writing – Original Draft, YI; Writing – Review & Editing, YI, and FE; Visualization, YI, YM; Funding Acquisition, FE.

## Classification

Biological Sciences, Neuroscience

