## Supplementary figures and legends. for "Divergent spatiotemporal integration of whole-field visual motion in medaka and zebrafish larvae"

### Supplementary information

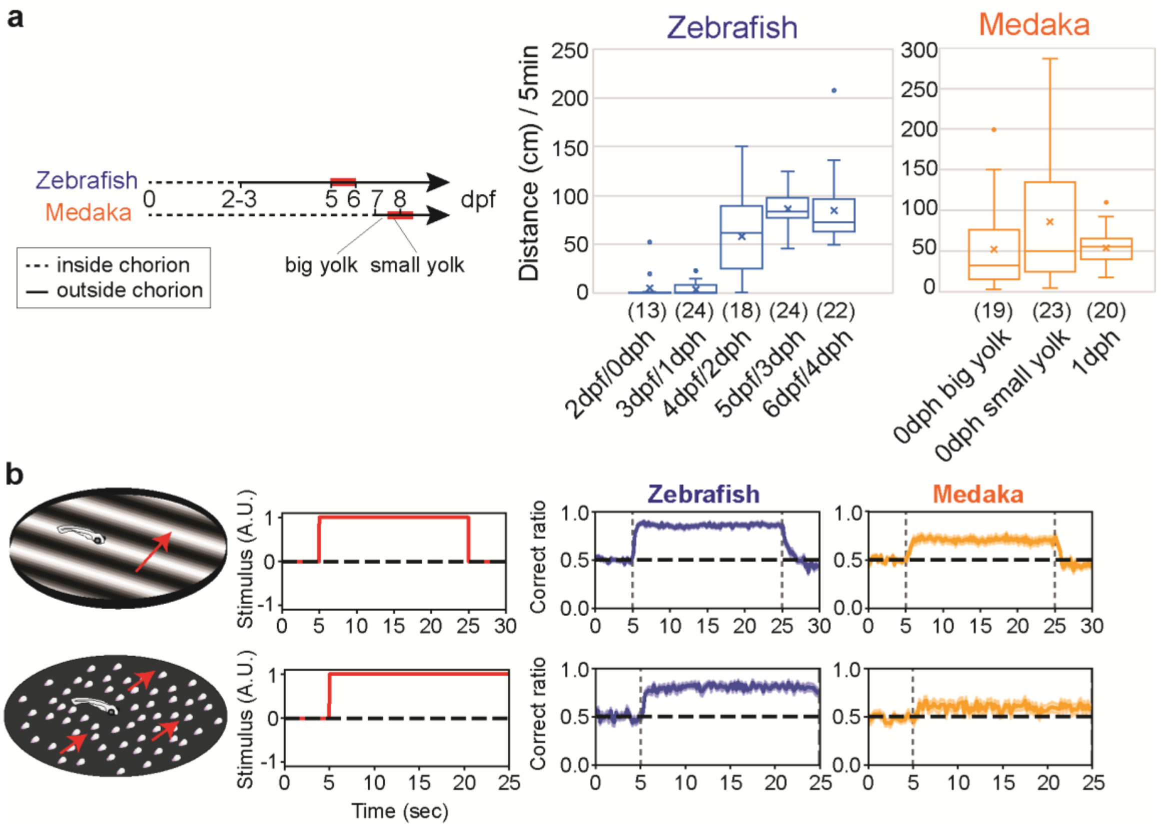


**Supplementary figure 1. Locomotion and OMR in zebrafish and medaka larvae**

**(a)** Developmental timeline of locomotion in zebrafish and medaka larvae. The two species exhibit different developmental rates: zebrafish larvae hatch at 2-3 days post-fertilization (dpf), while medaka larvae hatch at 7-8 dpf (left). However, zebrafish start regular swimming several days after hatching, whereas medaka initiate swimming immediately upon hatching (right panel). To match the developmental stage across species, we used 5-6 dpf zebrafish (3 days post-hatch, dph) and 7-8 dpf medaka (0-1 dph) for comparative analysis in this study. Red bars in the timeline indicate the sampling periods used for most experiments. Sample sizes for each measurement are indicated at the bottom of each graph in the right panel. **(b)** Quantification of optomotor response consistency. The "correct fraction" represents the proportion of swimming bouts oriented in the same direction as the visual motion stimulus. Average correct fractions (with shades of standard errors of means) for zebrafish (blue) and medaka (orange) in response to whole-field stimuli are shown for sine-grating stimuli (top row, zebrafish: n=53, medaka: n=42) and random dot motion stimuli (bottom row, zebrafish: n=21, medaka: n=32).

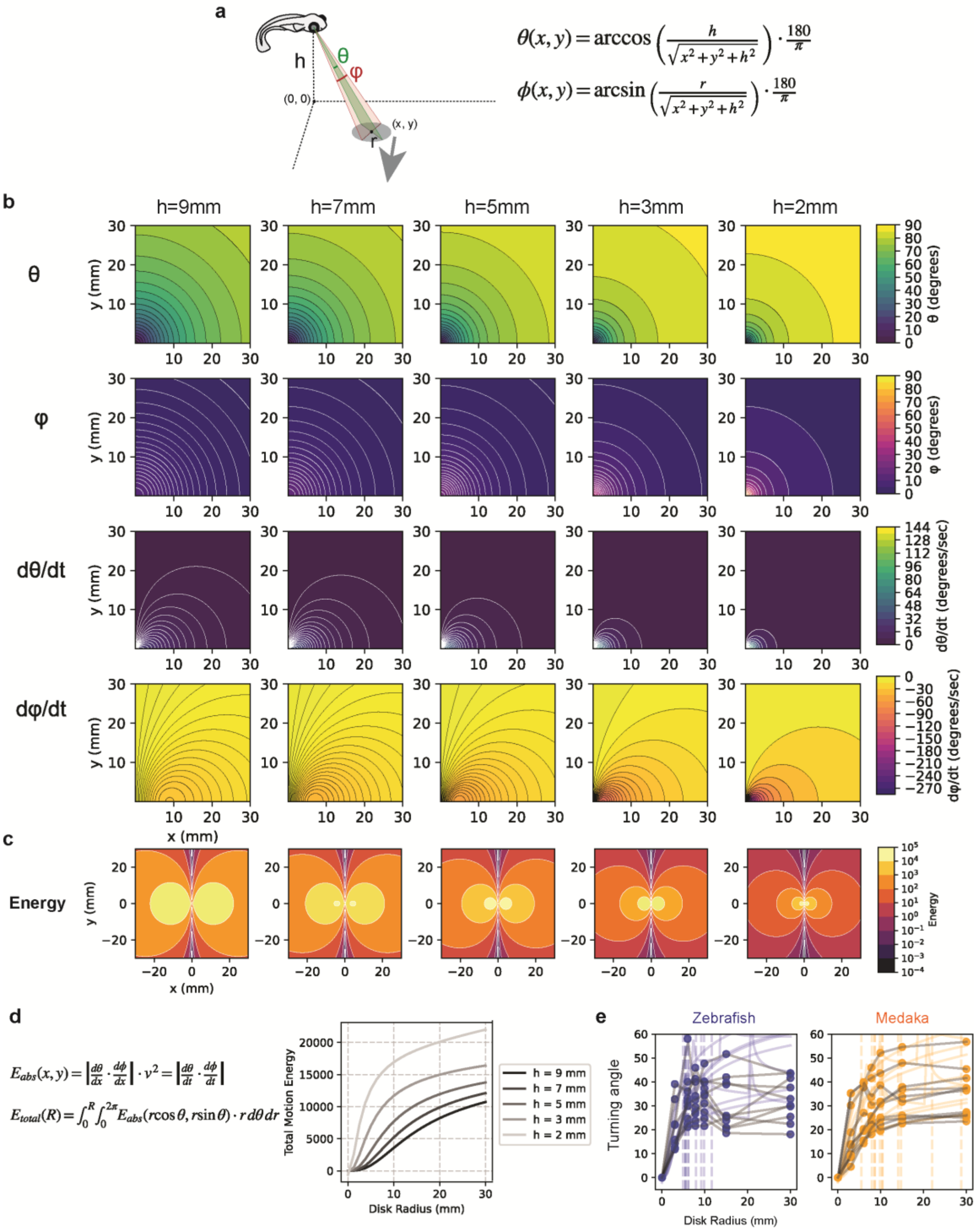


**Supplementary figure 2. Spatial integration of orthogonally moving dots**

**(a)** Angular metrics θ and φ defined for fish larvae at height h to calculate motion energy of dots at position (x, y) with radius r. Dots move left- or rightward, orthogonal to fish head direction (x-axis). **(b)** Angular parameters θ and φ, and their temporal derivatives dθ/dt and dφ/dt, vary dynamically with dot position and fish swimming height. **(c)** Motion energy generated by each dot varies dynamically with dot position and fish swimming height. **(d)** Total motion energy within the dot-filled disk increases with disk radius. Gray lines represent calculations at different fish swimming heights (the legends on the right). **(e)** Individual fish turning angles fitted to total motion energy function. Points represent experimental data (blue: zebrafish, orange: medaka); gray curves show fitted functions. Thresholds defined as disk radius where responses reach 95% of maximum turning angle (vertical dotted lines).

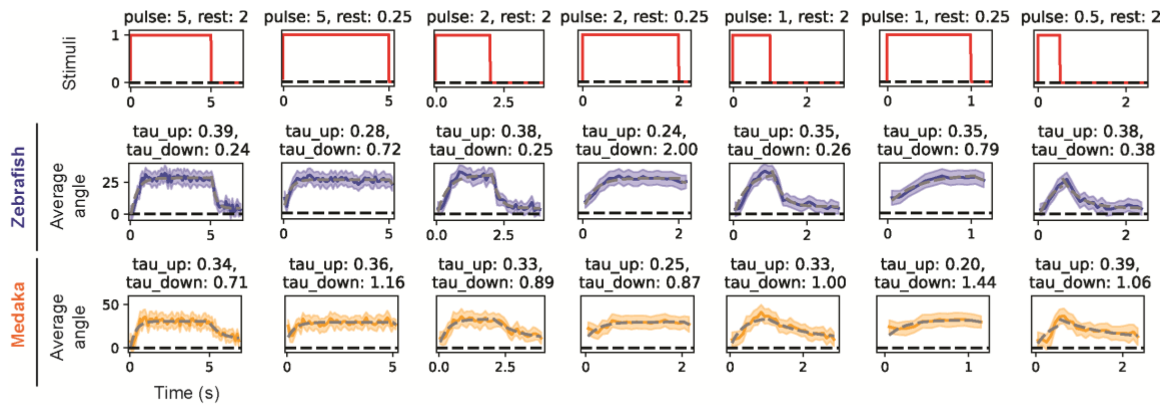


**Supplementary figure 3. Temporal dynamics of optomotor responses to pulse stimuli with varying durations.**

Behavioral responses to pulse stimuli with different pulse and rest durations. Following 5 s converging motion, repeated pulse-rest cycles of motion stimuli were presented until 25 s (protocol as in Figure 4a). Each panel shows average response during one pulse-rest cycle. Red traces: stimulus timing with labeled pulse and rest durations (unit: seconds) on top. Blue and orange traces: zebrafish (n=52) and medaka (n=21) responses. Shaded regions: mean ± SEM. Gray dashed lines: fitted exponential curves for estimating rise (tau_up) and decay (tau_down) time constants in seconds.

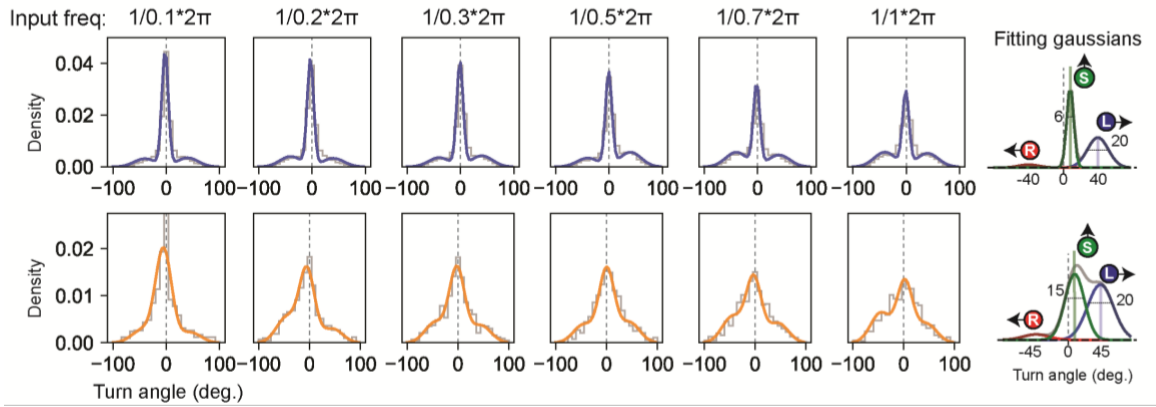


**Supplementary figure 4. Turn angle distributions fitted with three-component Gaussian models.**

Turn angle distributions of all turns recorded during alternating moving dot stimuli presentation. Gray lines: raw experimental data; colored lines: fitted three-component Gaussian curves (blue: zebrafish, orange: medaka). Each column represents responses at different stimulus frequencies. The three-component model captures distinct behavioral modes: large leftward turns, large rightward turns, and small directional adjustments.

### Supplementary movies legend

Please find the movies from the link:

https://drive.google.com/drive/folders/1KlxGbVVYm75BUdfHRhO_AAGTPc8rJYVs?usp=sharing

**S1 Movie. Sine-grating stimuli across the entire behavioral arena.**

Sine-grating visual stimuli were projected from below the dish. Fish position and head direction were recorded in real time, and visual stimuli were updated through online computation based on the fish's perspective. In the video, the blue dot indicates the position of the fish head and the line indicates the body axis. The direction of visual motion is always leftward from the fish's perspective. Visual stimuli cover the entire dish (radius of stimuli: 6 cm). Video playback speed matches the actual recording speed.

**S2 Movie. Sine-grating stimuli in a restricted area.**

Sine-grating visual stimuli were projected from below the dish. Fish position and head direction were recorded in real time, and visual stimuli were updated through online computation based on the fish's perspective. In the video, the blue dot indicates the position of the fish head and the line indicates the body axis. The direction of visual motion is always leftward from the fish's perspective. The radius of the visual stimulus is half the dish radius (radius of stimuli: 3 cm)., restricting the stimulus to the fish's immediate surroundings. Video playback speed matches the actual recording speed.

**S3. Movie Dot motion stimuli.**

Example video of dot motion stimuli. The lifetime of dots in this video is 100 ms
